# Age-dependent *Aedes* mosquito resistance profiling and mortality rate to repeated insecticides exposure in Western region, Saudi Arabia

**DOI:** 10.1101/2022.06.05.494159

**Authors:** Ashwaq M Al Nazawi, David Weetman

## Abstract

**Background:** Little is documented on *Aedes aegypti* age-dependent role on different resistance mechanisms to repeated insecticides exposures. The study examined the age-dependence of mortality rate and genetic resistance in two mechanistically pyrethroid resistant mosquito strains exposed once or repeatedly at different ages.

**Methods:** WHO bioassays and real time polymerase chain reaction (qPCR) were performed to ascertain their association between age-dependent exposures related mortality rate and single/repeated resistance in the Jeddah and Makkah. Candidate genes of interest (CYP9J7, CYP9J27, CYP9J26, AAEL006953, CYP9P450, AAEL006013) were assessment.

**Results:** Age dependent and exposure duration had a significant effect on the survival of the Jeddah and resistant Cayman. Our results showed that in a single exposure assays, age had no significant effect on mortality in the Cayman strain (χ2=2.76, df=1, *P*=0.097), but there was significantly increased mortality in the Jeddah strain younger age (χ2=5.46, df=1, *P*=0.02), but not statistically significant at older age. In the multiple exposure assay, GLiM analysis showed a significant strain, day and strain*day interaction indicating mortality rate is influenced by the strain or day (which also corresponds to age).The Jeddah strain showed generally lower survival,, there was a highly significant association of survival with repeated exposures in the Jeddah strain (χ2=43.6, df=1, *P*=4.1×10^E-11^) and the Cayman strain (χ2=12.5, df=1, *P*=0.0004). Mortality rate correlated statistically and significantly with the number of days of exposure in the Cayman strain (Spearman rank correlation ρ=-0.77, *P*=0.01), but in the Jeddah strain it was not statistically significant (ρ= -0.42, *P*=0.23). After repeated insecticide exposure, the AAEL006013 was statistically and significantly over-expressed compared to the control (*P*=0.03).

**Conclusion:** To the best of our knowledge, this is one of the first research on age and exposure linked genomic and bioassay on field *Ae. aegypti* in Jeddah, KSA. The study showed that repeated exposure to pyrethroids reduced the *Aedes* mosquito population mortality rate. This suggests that there is indeed increasing age-dependent resistance or survival with multiple exposure high-doses of same or repeated insecticide, thus indicating the need to rethink on integrated vector control policy and interventions and technical assistance in the Kingdom.

## Introduction

Dengue is a viral disease transmitted by the bite of *Aedes* mosquitoes, mainly *Ae. aegypti*, threatening economies and human health in most tropical regions of the world. In Saudi Arabia, dengue has remained endemic since the first case was reported in 1994 in Jeddah [1], with subsequent spread to Makkah and Jizan. Insecticide-based control of *Ae. aegypti* remains the main dengue control option in Saudi Arabia as currently there is no preventative or curative medication, and the approved vaccine [2], is not yet available in the Middle Eastern region.

To monitor insecticide resistance, the World Health Organization (WHO) recommends the use of 3 to 5-day-old, non-blood-fed female mosquitoes, which have not been previously exposed to insecticide in diagnostic dose bioassays[3]. This standardisation aims to reduce confounding variables, which can greatly impact test results[4], and to facilitate comparison among different tests, permitting resistance surveillance and monitoring over time. The standard WHO bioassays showed that both *Aedes* mosquito populations from Jeddah and Makkah were resistant to permethrin, deltamethrin, and bendiocarb[5]. Resistance target site *kdr* mutation and P450-based metabolic mechanisms were identified in the Makkah and Jeddah strains [5]. However, no *Ace-1* mutation has been studied yet. At very extreme weather and high temperature perhaps for females at the upper end of day 3 to 5, dengue transmission to humans is unlikely to occur prior to 8 days post-emergence, with 14 or more days probably more typical[6]. This period encompasses females mating, taking an infectious blood meal and surviving the extrinsic incubation period (EIP) during which the virus replicates and migrates to the salivary glands for transmission during the next blood feeding. Nevertheless, in addition to standardisation of assays, testing a predominantly pre-transmission age group may still be the most disease control-relevant option if (i) survival of a single insecticide exposure declines with age, or, if this is not the case, that (ii) repeated exposure experienced by older mosquitoes has cumulative effects due to selective pressure exerted by the active metabolite and residual insecticides derivatives which result in their reduced survival or emergence of resistance. To determine whether these scenarios apply, it is important to study the age- and exposure-specific survivorship profiles of mosquitoes to allow prediction of effects of control on disease transmission. Previous studies on *Aedes* and *Anopheles* have shown that susceptibility to insecticides increases with chronological age [7-11]. For example, 3 day-old *Ae. aegypti* females were significantly more tolerant to 4% DDT (mortality ∼<10%) and 0.05% deltamethrin (mortality ∼60%) in WHO bioassays after 24h, than 14-day-old females(DDT mortality ∼40%; deltamethrin mortality ∼<80%) [7]. However, in spite reported age-dependent pyrethroid survivorship trend in the literature[4], it is unclear if this may depend on resistance mechanisms expressed by the strains examined, e.g. metabolic enzymes versus target site mutations and metabolic (P450 enzyme expression)-based resistance, both of which are common in Jeddah and Cayman *Ae. aegypti* strains, [12]. Poupardin and colleagues demonstrated that repeated exposure of *Ae. aegypti* to xenobiotics and insecticides leads to induction of CYPs and GSTs [13].

The study investigated the patterns of age-dependent *Aedes* mosquito and multiple pyrethroid exposure linked resistance from single versus repeated exposures over time.

## Methods

### Mosquito Strains

Field-collected *Ae. aegypti* mosquitoes from two dengue endemic areas in Makkah (N21°40’7.70, E39°86’3.19) and Jeddah (N21°60’3.97, E39°27’2.49). The Saudi Arabian *Ae*.*aegypti* strain was originated from larval collected from several breeding sites within Jeddah and Makkah. The larvae were reared as described by Alnazawi et al.[5]. Cayman, a resistant lab strain [14], was chosen as a comparator because unlike the Jeddah strain, it showed no evidence for PBO synergy of deltamethrin resistance[4, 5], with pyrethroid resistance apparently primarily dependent on *kdr* mutations (V410L, V1016I, V1534C)[15]. For quantitative PCR assays of candidate resistance genes, the standard susceptible strains New Orleans, Rockefeller and Liverpool strains were used as reference. All strains were raised under the same conditions [5].

### Bioassays

Jeddah female field *Aedes* mosquito first generation (F0) and Cayman resistant mosquitoes of different ages were exposed to WHO deltamethrin papers to test the hypothesis that susceptibility increases with age. Jeddah field is used for bioassays and *Ace-1* sequencing target site, and molecular work, whereas Makkah used for target site. In the single exposure assay, pools of 25 mosquitoes per replicate of ages between 5 and 14 days (separately) were exposed to deltamethrin control paper for 1h. After the exposure, females were transferred to recovery tubes and provided with 10% sucrose. In the multiple exposure assay, and unfed female mosquitoes (five days old) Cayman and F0 Jeddah mosquitoes were exposed to 0.05% deltamethrin for 1h. After the 24h recovery period, all survivors were re-exposed to the same deltamethrin dosage. The exposure was continued every day until the remaining mosquitoes were 14 days old to test the hypothesis that repeated exposure would increase the mortality rate. Mortality was recorded every day. All survivors from single and multiple exposure were later preserved in RNA and stored at -20°C before qRT-PCR analysis was conducted.

### Statistical analysis

The effects relating to the age of the strains were analysed using generalised linear models with binomial link functions in SPSS version 24. Error bars represent 95% confidence intervals. The cumulative mortality analysis was performed in GraphPad prism7.

## Results

### Age dependence and single or multiple-exposure deltamethrin-induced mortality

The GLiM analysis showed a significant association of mortality with age in the Jeddah strain (χ2=14.66, df=1, *P*=0.000129), but not in the Cayman strain (χ2=1.619, df=1, *P*=0.203) **(Table 1)**.

**Table 1.** Generalized Linear Model for the effects of strain and age on deltamethrin-induced mortality of *Ae. aegypti* females.

| Source | Wald $\chi^2$ | df | Probability |
| --- | --- | --- | --- |
| (Intercept) | 15.4 | 1 | 0.000 |
| Strain | 0.423 | 1 | 0.516 |
| Day | 14.54 | 1 | 0.000 |
| Strain * day | 5.14 | 1 | 0.023 |

Our results showed that in a single exposure assay, age had no significant effect on mortality in the Cayman strain (χ2=2.76, df=1, *P*=0.097), but there was significantly increased mortality in the Jeddah strain (χ2=5.46, df=1, *P*=0.02) **(Fig.1)**. Although there was a significant difference in mortality in respect of the age and degree of exposure in the Jeddah strain, the impact of age on mortality was not different until the oldest age (14 days). Therefore, mortality-age association was not a simple linear relationship.

In the multiple exposure assay, GLiM analysis showed a significant strain, day and strain*day interaction indicating mortality rate is influenced by the strain or day (which also corresponds to age) **(Table 2)**.The Jeddah strain showed generally lower survival, there was a highly significant association of survival with repeated exposures in the Jeddah strain (χ2=43.6, df=1, *P*=4.1×10^E-11^) and the Cayman strain (χ2=12.5, df=1, *P*=0.0004).

**Fig 1.**
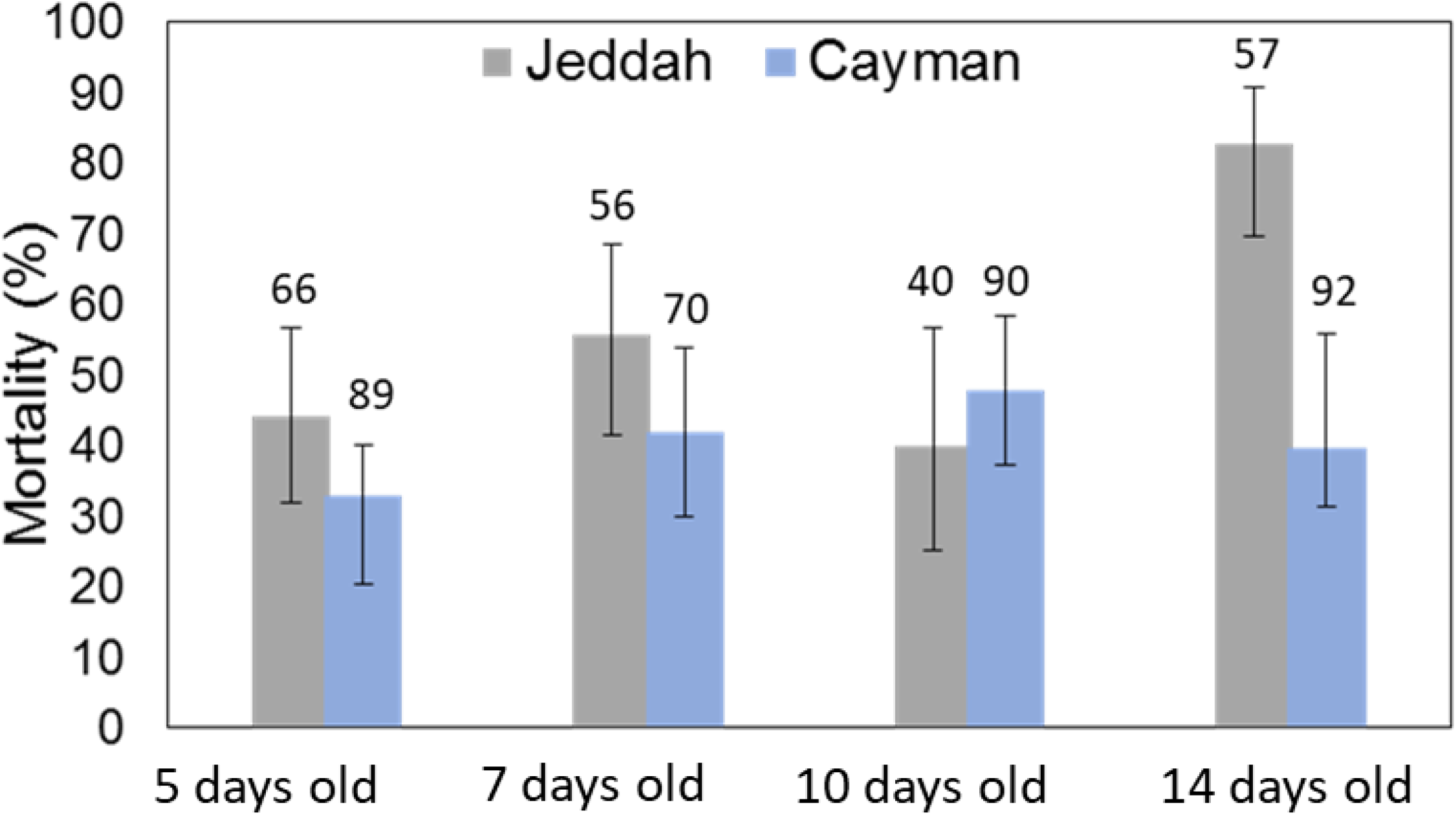
Single exposure of Cayman and Jeddah mosquitoes to deltamethrin for 1h at age 5, 7, 10 and 14 days. The number of *Ae. aegypti* mosquitoes assayed is presented above each bar.

**Table 2.** Generalized Linear Model for effects of multiple exposure to deltamethrin and strain on mortality of *Ae. aegypti* females.

| Source | Wald $\chi^2$ | df | Probability |
| --- | --- | --- | --- |
| (Intercept) | 36.929 | 1 | 1.2252E-9 |
| strain | 10.274 | 1 | 0.001349 |
| Day | 16.383 | 1 | 0.000052 |
| strain * day | 5.572 | 1 | 0.018 |

Interestingly, high mortality was observed at the beginning of insecticide exposure which progressively declined as shown by the flattening of the cumulative mortality curve **(Fig.2)**. The mortality rate in the Jeddah and Cayman strains reduced from 43.7% and 15.1% in day 1 to 0% and 6.3% in day 10 respectively. Mortality rate correlated statistically and significantly with the number of days of exposure in the Cayman strain (Spearman rank correlation ρ=-0.77, *P*=0.01), but in the Jeddah strain it was not statistically significant (ρ= -0.42, *P*=0.23).

**Fig 2.**
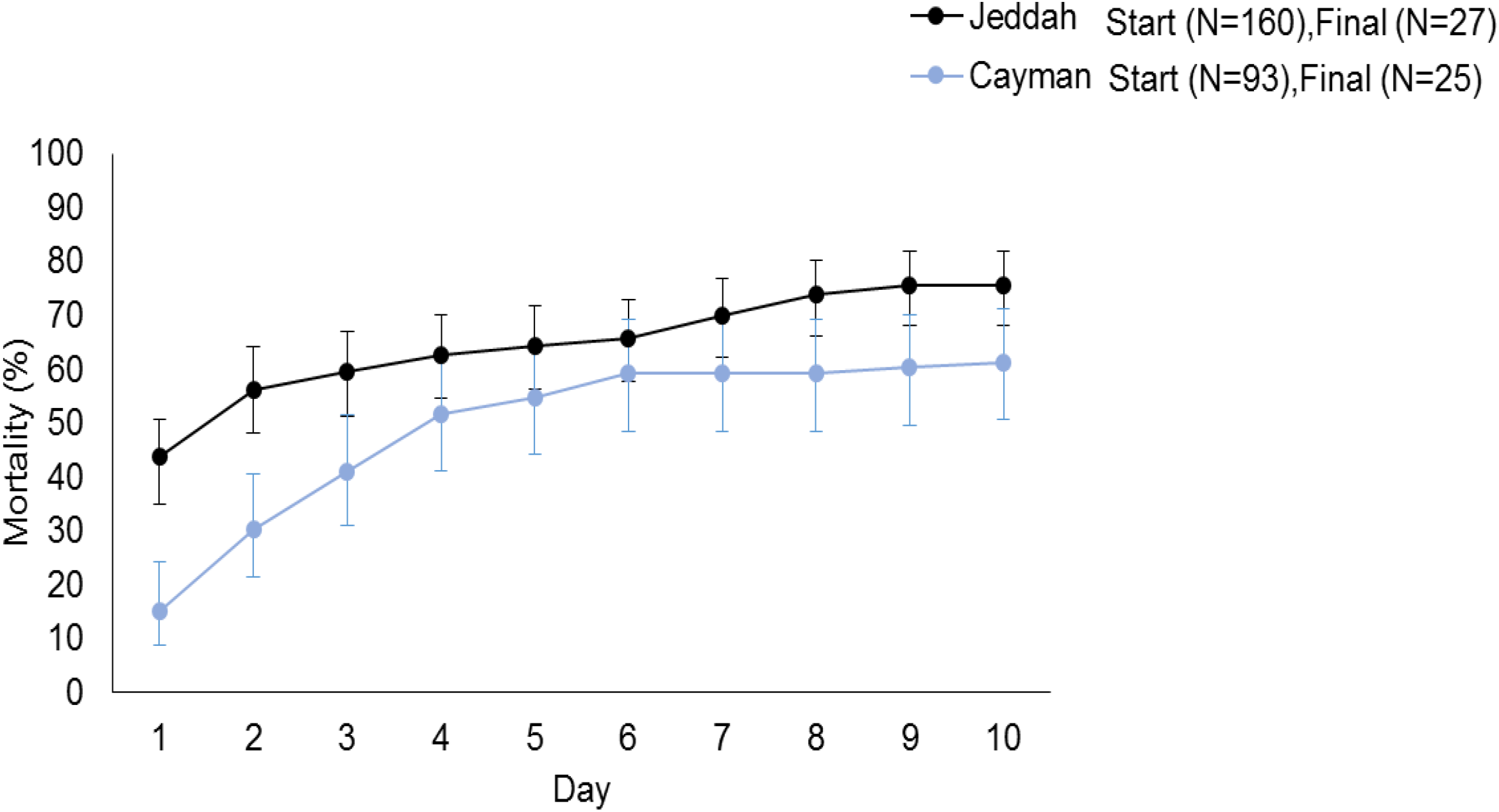
Cumulative mortality for each strain on different days. The x-axis represents the number of mosquitoes at the beginning of the experiment and the y-axis is the number of mosquitoes alive by Day 10.

### Effect of single and repeated deltamethrin exposure on mortality of 10 old mosquitoes

Ten-day old Cayman and Jeddah strains had lower mortality in the group that had been repeatedly exposed (6 times/1h) to deltamethrin compared to the group exposed only once to deltamethrin illustrated in a previous study [5] for either 1h, 6h or 8h (**Fig.3)** because of the diminishing rate across exposures.

**Fig 3.**
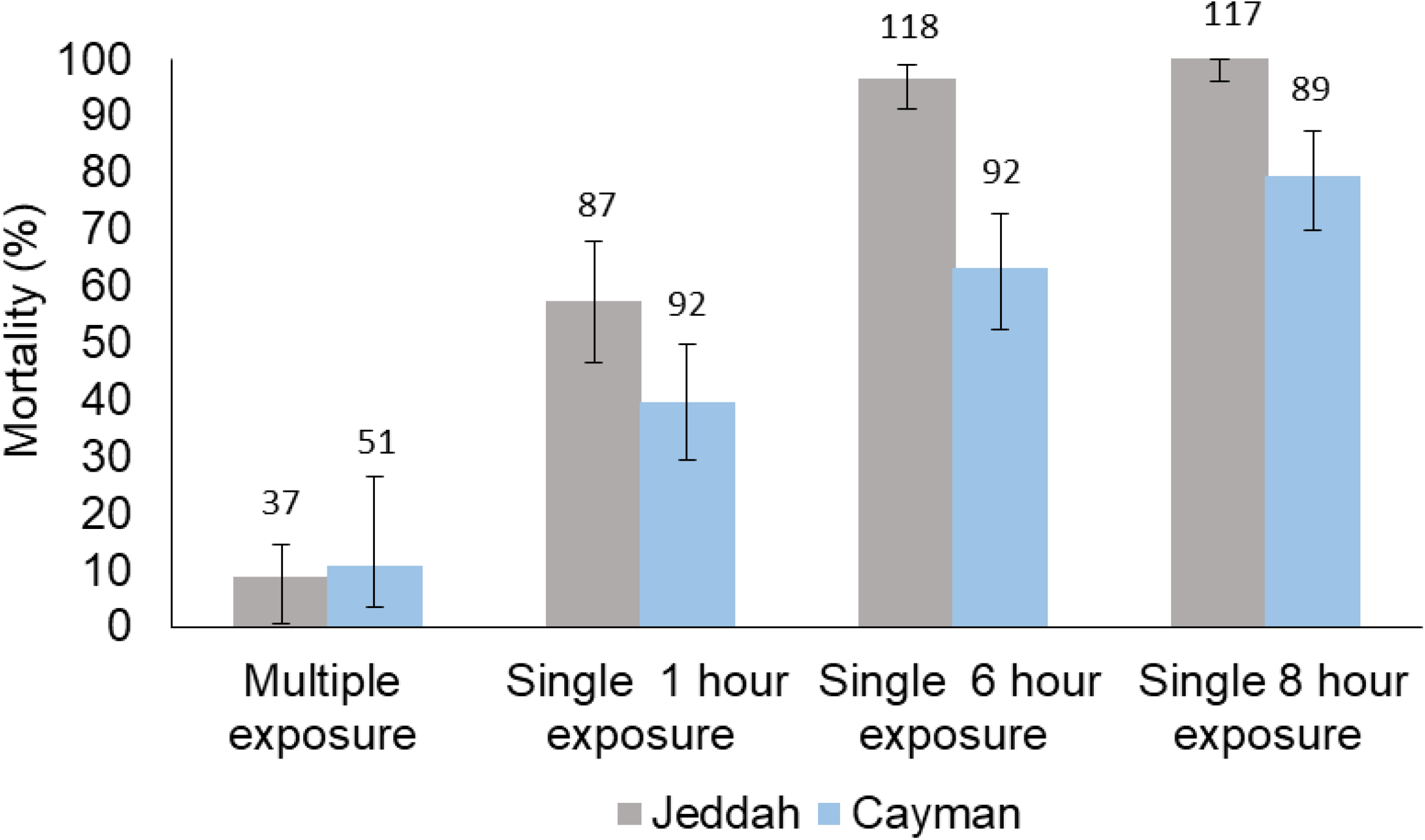
A composite figure comparing the effect of different deltamethrin exposure durations on mortality of ten-day old females. In grey and blue is 10 day old Jeddah and Cayman strain, respectively. Error bars represent 95% confidence intervals. The number of *Ae. aegypti* mosquitoes assayed is presented above each bar.

### Sequencing of the Acetylcholinesterase 1 (*Ace-1*) gene

The *Ace-1* gene was successful amplified in 10 individuals and 5 pools of 10 mosquitoes from the Jeddah and Makkah field *Aedes* mosquito populations studied. Neither the G119S, nor any other mutation was identified in the *Ace-1* gene fragments sequenced from the Saudi strains **(Fig.4)**.

**Fig 4.**
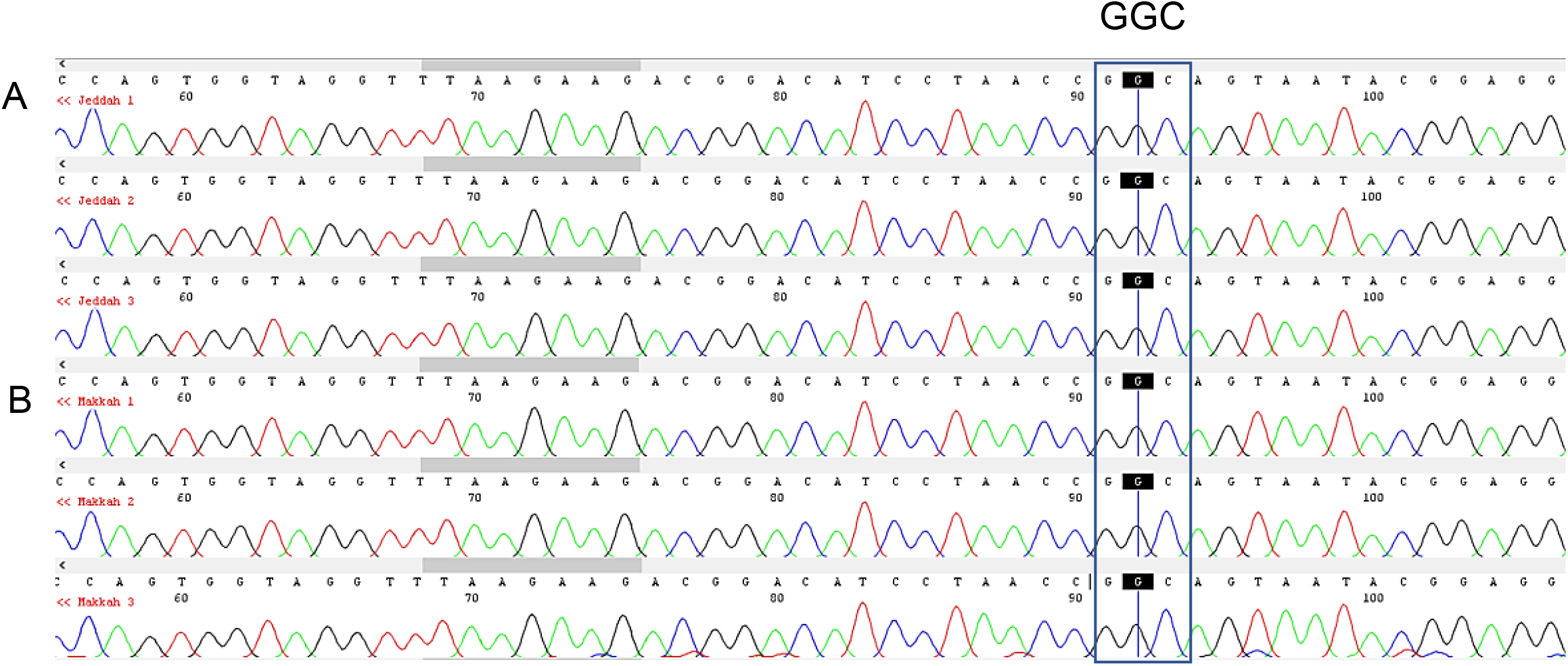
Chromatograms of nucleotide sequence from Codon code aligner software illustrating no mutation at position 119. Example of samples A) Jeddah, B) Makkah. GGC is a wild-type codon.

### Gene expression analysis

#### Age-dependent expression of metabolic genes

Our findings on the expression level of each gene in mosquitoes aged 3, 5, 10 and 14 days old relative to the susceptible strains revealed that there was no significant difference between the different ages **(Table 3)**.

**Table 3.** Gene expression in strain aged 3, 5, 10 and 14 days relative to the three susceptible strains in Jeddah using one-way ANOVA.

| Gene | F test | df | *P* value |
| --- | --- | --- | --- |
| CYP9J7 | 2.8 | 3 | 0.06 |
| CYP9J27 | 0.58 | 3 | 0.6 |
| CYP9J26 | 1.4 | 3 | 0.3 |
| AAEL006953 | 0.49 | 3 | 0.05 |

Relative-fold changes compared to three susceptible strains of CYP9J26, Rockefeller were 250, 150, 100 times higher susceptibility at Day 3, Day 5 and Day 10 old respectively. Through qPCR analysis, the differential expression of CYP9J7 and AAEL006953 genes were statistically and significantly overexpressed in the pyrethroid resistant Jeddah strain from Saudi (**Fig.5)**.

**Fig 5.**
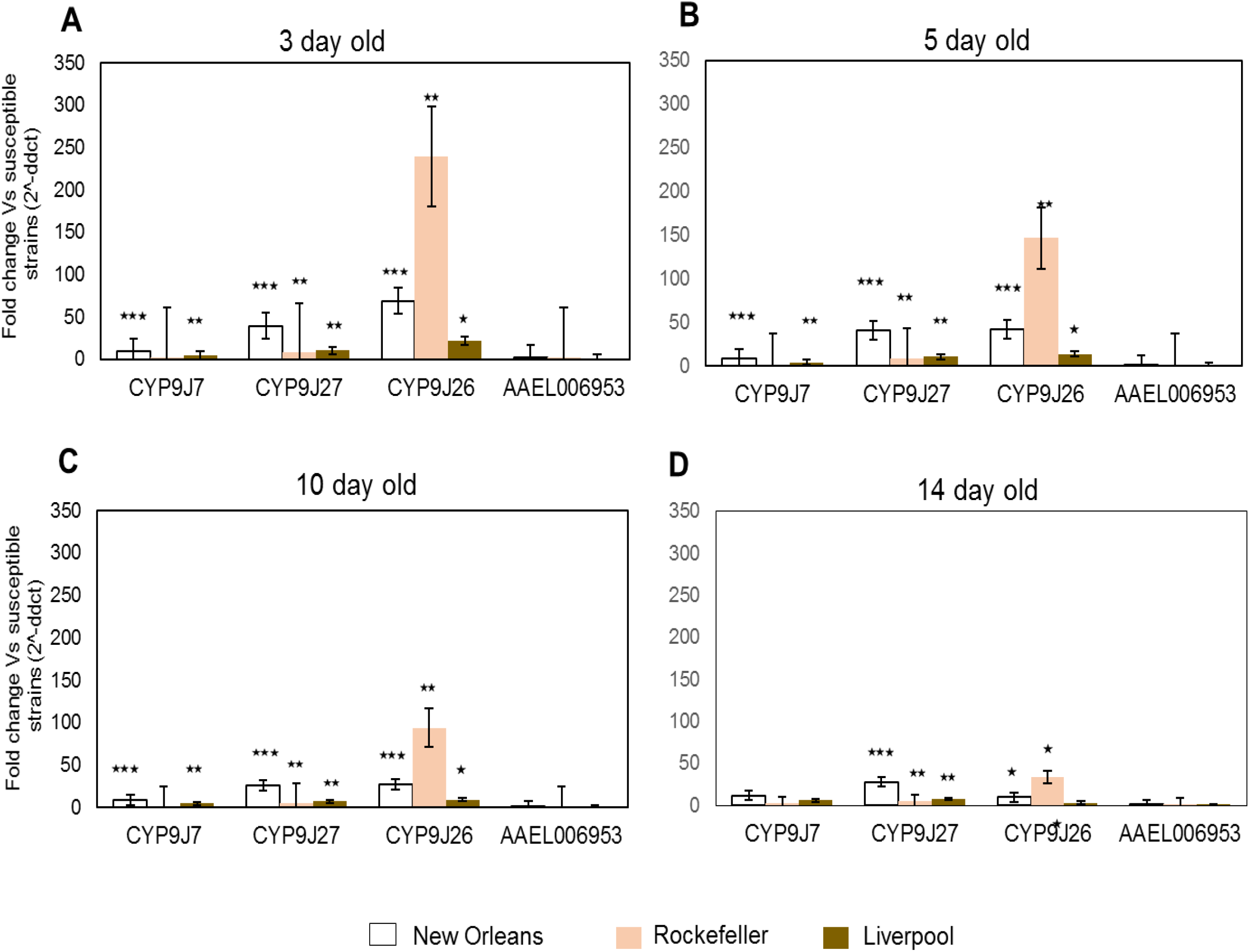
Quantitative PCR analysis of candidate genes on age-dependant mortality. Relative-fold changes compared to three susceptible strains (New Orleans, Rockefeller, Liverpool) are shown following normalisation to two endogenous reference genes. Error bars represent Standard Error (±SE). (Two-tailed *t* test, \**P*< 0.05, \*\**P*< 0.01 and \*\*\**P*< 0.001).

#### Effect of multiple or repeated insecticide exposure on gene expression

Our results showed that lower mortality to deltamethrin was observed when mosquitoes were repeatedly exposed to it compared to unexposed mosquitoes. In those repeatedly exposed to deltamethrin, the expression level of the following tested genes CYP9J7, AAEL014614-RA (CYP9P450) and AAEL006953-RA was higher (but not significantly) in comparison to controls but no difference was seen in CYP9J27 and CYP9J26 **(Fig. 6)**. AAEL006013 was statistically and significantly overexpressed in repeated exposure compared to the control to deltamethrin (*P*=0.03) **(Table 4)**.

**Table 4.** The difference of gene expression between repeated exposure to deltamethrin and control relative to the three susceptible strains using a *t* test.

| Genes | Repeated non-exposure compared to repeated exposure |  |  |  |
| --- | --- | --- | --- | --- |
|  | F test | t value | df | *P* value |
| CYP9J7 | 0.384 | 1.218 | 16 | 0.241 |
| CYP9P450 | 0.728 | 0.96 | 16 | 0.351 |
| CYP9J26 | 0.909 | 0.546 | 16 | 0.592 |
| AAEL006013 | 0.869 | 2.378 | 16 | 0.030 |
| CYP9J27 | 0.823 | 1.619 | 16 | 0.125 |
| AAEL006953 | 0.488 | 0.193 | 16 | 0.848 |

**Fig 6.**
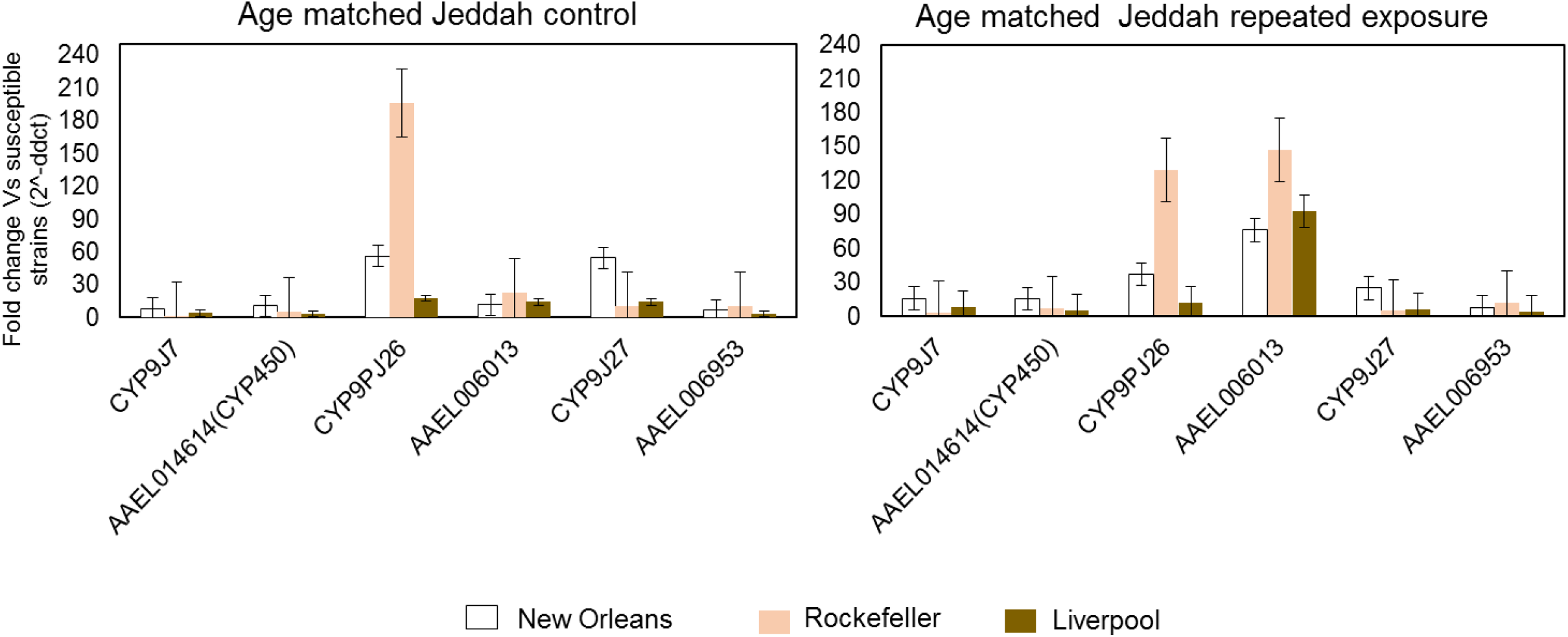
Quantitative PCR analysis of candidate genes following multiple exposure. Relative-fold changes compared to three susceptible strains (New Orleans, Rockefeller, Liverpool) are shown following normalisation to two endogenous reference genes. Error bars represent Standard Error (±SE). Age matched Jeddah control and exposure were compared.

In age matched Jeddah control CYP9PJ26 Rockefeller strain was 210 times fold changes, whereas age matched Jeddah repeated exposure showed the overexpression to deltamethrin of 120 and 50 fold increase n CYP9PJ26 and AAEL006013 genes **(Fig. 6)**.

## Discussion

Data gaps on *Aedes* mosquito susceptibility to candidate insecticides is a limiting factor for the success of control programmes, where these programmes have often been implemented without full information on the resistance risk posed by a given control agent in the field. Therefore, the current study was conducted to assess both the effect of age and multiple insecticide exposure encounters over time and the mechanisms.

Continuous and repeated vector control programs based on the use of insecticide could be linked with increasing selective pressure to same insecticide due to suboptimal dose in the environment and increasing emergence due selective pressure on target genes site and disruption of mechanism of action by low doses of pyrethroid or deltamethrin. Advisable, evidence based genomic and bioassay is capital is directing vector control programs mainly the use of insecticide to target sites to reduce *Aedes* mosquito vector burden.

Moreover, *Aedes* longevity is a key factor for disease transmission in disease vectors as prolonged survival of vectors in the wild increases the chances of their becoming infected, and of a successful completion of the extrinsic incubation period of the pathogens allowing subsequent transmission in later feeding events. In the study on *Aedes* and other species conducted by Al Nazawi et al.(3), age has been negatively associated with insecticide exposure survival even in resistant populations [6-8, 16-18]. For example, in the case of unfed four-day-old *An. funestus*, survival was significantly higher than older (10-day-old) mosquitoes after 24h post exposure to 0.1% lambda-cyhalothrin [18]. Rajatileka et al. [7] observed that unfed young (three-day-old) *An. gambiae* lab strains from Zanzibar-Tanzania, Kisumu-Kenya, and Akron-Benin and *Ae. aegypti* from Merida in Mexico and Ho Chi Minh in Vietnam reported survival significantly more after 24h post-exposure to DDT, bendiocarb and deltamethrin compared to their 14-day-old unfed counterparts [7].

In this study, ten-day-old females *Ae. aegypt*i were significantly more susceptible to deltamethrin than those which were three to five days old in a standard WHO susceptibility test in Jeddah[5]. However, age-dependent increased susceptibility to deltamethrin was not observed in the Cayman strain, thus indicating that the strain genetic is not universal across all mosquito populations [5].Most arboviruses require an extrinsic incubation period in the vector that ranges from between 7 to 14 days [19]; over this susceptibility period to pyrethroids increased significantly in the Jeddah strain (38.6% increase mortality in 14-day-old mosquitoes compared to those which were five days old). Therefore, if pyrethroids are used to control *Aedes*, and they are applied correctly (with respect to dose, timing, frequency,), increased susceptibility with age could help to reduce arbovirus transmission. We observed a reduction in mortality in 14-day-old females *Aedes* that had been repeatedly exposed to deltamethrin every 24h compared to a three-day-old cohort when first exposed. This result contrasts with those in the age-dependent mortality study. Repeated exposure every 24h may have led to either selection of the most genetically resistant individuals and/or induction of detoxification genes. Irrespective of the mechanism, given the importance of adult longevity for disease transmission [20], the finding that repeated exposure o insecticide can upset the normal age-dependent susceptibility relationship of *Aedes* mosquito population. This is a concern for pyrethroid-based control in integrated vector management in Western regions, and it will be important to rethink and improve innovative biocontrol measures and interventions in KSA.

In qRT-PCR on age dependent expression of genes, revealed that there was no significant difference between the different ages the expression level of each gene in mosquitoes aged 3, 5, 10 and 14 days old relative to the susceptible strains **(Table 3)**. Rockefeller susceptible CYP9J26 strains were 250, 150, 100 times more susceptible at Day 3, Day 5 and Day 10 old respectively. Through qPCR analysis, showed differential expression of CYP9J7 and AAEL006953 genes were statistically and significantly overexpressed in the pyrethroid resistant Jeddah strain from Saudi. It can be interpreted that downregulation of metabolic genes was associated with increasing selective pressure due to repeated insecticides exposure or emergence of insecticide resistance may explain why decreasing/increasing susceptibility with mosquito age [16, 21]. For instance, although not significant, the expression levels of the cytochrome P450 CYP9J26 in the Jeddah strain, which has been shown to metabolise deltamethrin *in vitro*[22] and is frequently overexpressed in pyrethroid resistant populations [12], decreased from a fold change of 68.6 relative to three-day-olds in the New Orleans strain, 42 in those which are five days old, 27 in those which are ten days old and 9.6 in 14-day-old mosquitoes when assessed by qRT-PCR **(Fig.5)**. A similar reduction in the expression level of this gene in respect of the Rockefeller and Liverpool strains was observed **(Fig.5)**.

Lower mortality was reported to deltamethrin when mosquitoes were repeatedly exposed to it compared to unexposed mosquitoes. Repeatedly exposure to deltamethrin showed increasing expression level of genes CYP9J7, AAEL014614-RA (CYP9P450) and AAEL006953-RA (but not significantly) in comparison to controls **(Fig.6)**. These results are consistent with previous reports (4,5,9,15,21). Interestingly, AAEL006013 gene was statistically and significantly overexpressed in repeated exposure compared to the control to deltamethrin (P=0.03) **(Table 4)**. His finding indicates that gene-related selective pressure could be linked with targeted mechanisms of action of the insecticide to single or repeated exposure.

In age matched Jeddah control CYP9PJ26 Rockefeller strain was 210 times fold changes, whereas age matched Jeddah repeated exposure showed the overexpression to deltamethrin of 120 and 50 fold increase n CYP9PJ26 and AAEL006013 genes Regulation of this gene and CYP9J27 is highly likely to be age-dependent rather than induced by insecticides since the genes were expressed in low levels in older mosquitoes that had been repeatedly exposed to insecticides. Other studies have reported similar findings where metabolic genes such as GSTE2, GSTE1, CYP6P3, CYP6P4, CYP6Z3, CYP6M2 and COEAE1A have consistently been found overexpressed in resistant *Aedes* mosquito population were found to be stage or developmentally regulated [23-26]. Multiple or repeated exposure to pyrethroids diminished the mortality rate, suggesting that there are indeed higher resistance in *Aedes* mosquito population that can survive either high-level exposure or multiple exposure.

Data on the efficacy of control intervention against *Aedes* is limited. Therefore, it is difficult to ascertain whether or not insecticides will remain effective in controlling transmission of arboviruses by *Aedes* populations which are highly resistant to insecticides [12].

## Conclusion

To the best of our knowledge, this is the first genomic and bioassay analyses research study on field *Aedes* mosquito age and repeated insecticide exposure; it revealed that repeated exposure to pyrethroid or Deltamethrin diminished the mortality rate at younger age or increasing resistance emergence in *Aedes* mosquito population. Age and insecticide exposure are indeed increasing age-dependent resistance or survival with multiple exposure high-dose of same insecticide. This is an important finding in rethinking or reformulating vector control strategies and public health implications in western regions in Saudi Arabia. Though, Dengue remains a persistent public health burden and threats in western regions (mainly Jeddah and Makkah) of Saudi Arabia.

## Abbreviations

PBO: Piperonyl butoxide;
EIP: extrinsic incubation period;
Ace-1: Acetylcholinesterase;
CYP450s: cytochrome P450s;
GST:GSTs: glutathione-s-transferases;
Kdr: knockdown resistance;
DDT: dichlorodiphenyltrichloroethane;
WHO: World Health Organization;
qRT-PCR: quantitative reverse transcription polymerase chain reaction;
KSA: King Saudi Arabia ;
GLiM: generalised linear model;
SPSS: Statistical Package for the Social Sciences.

## Acknowledgements

Authors are grateful to all the municipal staff (administrators and laboratory technicians in the vector control programme) of Makkah and Jeddah in the Saudi Arabia. We also thank all staff in the vector control centre of Makkah, who participated in the mosquito collection.

## Data availability

Data are supplied in manuscript tables or figures

## Authors’ contributions

AMA-N collected the field samples, performed the insectary bioassays and molecular analyses, analysed data and drafted the manuscript. DW conceived and designed the experiments, drafted the manuscript and analysed data.

## Consent for publication

Not applicable.

## Ethics approval and consent to participate

Not applicable.

